# Nutritional Ketosis Attenuates Sucrose Bingeing-Induced Behavioral Deficits by Improving Synaptic Plasticity and Anti-Inflammatory Signalling in the Prefrontal Cortex

**DOI:** 10.1101/2025.05.07.651641

**Authors:** Chitralekha Gusain, Mohit Kumar

**Affiliations:** National Agri-Food and Biomanufacturing Institute (BRIC-NABI), S.A.S Nagar, Punjab, 140306, India; Regional Centre for Biotechnology, Faridabad, Haryana, 160014, India; Adjunct faculty, Regional Centre for Biotechnology, Faridabad, Haryana, 160014, India

**Keywords:** Anxiety, Binge eating disorder, Blood-brain barrier, Neuroinflammation, Nutritional ketosis, Sex differences

## Abstract

Binge eating, one of the defining characteristics of binge eating disorder, has been linked to poor health span. Animals, like humans, selectively binge on highly palatable foods, implying that the binge-like behavior is motivated by hedonic rather than metabolic signals. Given the association between reward processing and food intake, this study explored sex-specific underlying biological mechanisms, including synaptic, neuroimmune and neurometabolic correlates of binge-like sucrose drinking, compulsivity and anxiety-like behaviors, utilizing a preclinical model of hedonic sucrose drinking. Throughout the experiment, male and female mice binged on a sweet solution when given food *ad libitum* with free access and a choice between water and a 10% (w/v) sucrose solution, with females consuming more sucrose. The sucrose intake was positively correlated with transcription of genes for dopamine receptors (*Drd1, Drd2*) in the PFC of male and female mice. Sucrose-bingeing increased PFC neuroinflammation, concomitantly increasing region-specific blood-brain barrier permeability in males. Sucrose bingeing elevated the transcription of glucose metabolism genes (*Slc2a3, Glo1*) while inhibiting ketone oxidation pathway genes (*Slc16a1, Oxct1, Acat1*) in the PFC of males and females. Nutritional ketosis attenuated sucrose bingeing, compulsivity and anxiety-like behaviors in sucrose-dependent male and female mice by suppressing the transcription of reward-related genes (*Drd1, Drd2*) while promoting an anti-inflammatory neuroimmune microenvironment in the PFC of male and female mice in a sex-dependent manner. Overall, this study identified synaptic, neuroimmune and neurometabolic mechanisms as novel druggable targets and nutritional ketosis as a potential therapy for diseases where binge-like eating, compulsivity and anxiety-like behaviors are the main symptoms, such as binge eating disorder.

**Graphical Abstract:** 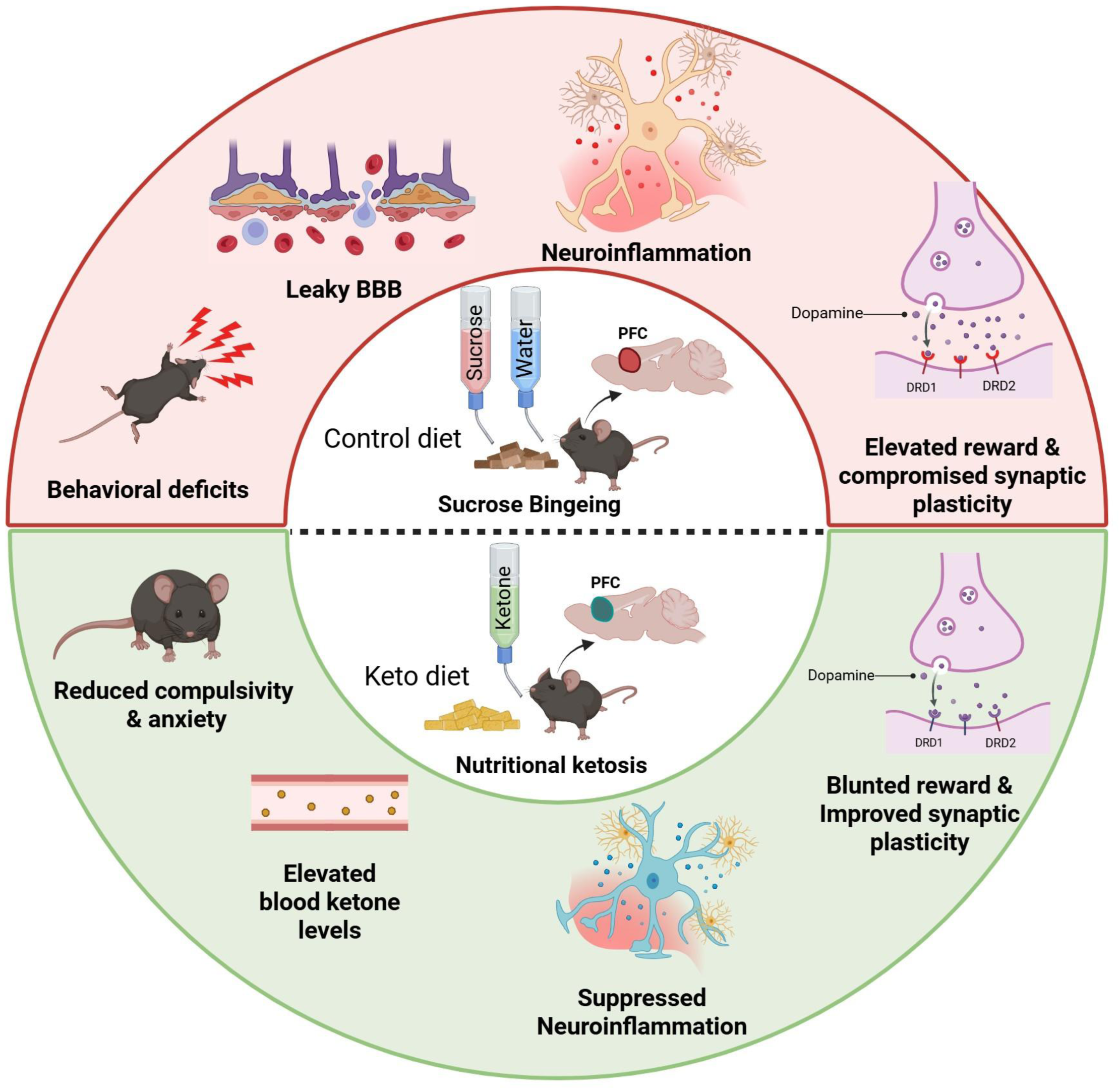

## 1. Introduction

Binge Eating Disorder (BED) is a serious but treatable mental health problem that is now classified as an autonomous disorder in the DSM-5(1). BED is the most prevalent eating disorder among all other eating disorders, affecting 2–5% of the adult population and is more common in females than males(2, 3). BED is defined by regular episodes of eating substantial amounts of food (typically within 2 hours), frequently quickly and to the point of discomfort. Unlike other eating disorders, such as bulimia nervosa, patients with BED do not engage in compensatory behaviors such as vomiting, fasting, or excessive exercise following binge eating. Although binge eating has been associated with substantial medical problems, including an elevated risk of psychiatric disorders(4), therapeutic options for BED are limited.

Globally, dietary or added refined sugar consumption has been increasing exponentially. Most of this comes from ultra-processed sugary beverages such as soda, energy drinks, fruit juices, sweetened coffee and tea, syrups, and sports drinks(5). Sugars are rewarding and can activate neural pathways and mechanisms linked to addiction, changing eating behavior from homeostatic to hedonic overeating(6). Sugar bingeing increases the risk of metabolic diseases, such as type 2 diabetes and obesity(5), neurological conditions such as neuroinflammation(7, 8) and neurovascular dysfunction(7) and psychiatric conditions such as cognitive decline(9, 10), motor learning impairment(11), depression and anxiety(12, 13). Sugar bingeing also causes brain insulin resistance in glial cells of *Drosophila melanogaster*, which impairs glial phagocytic function through compromised energy metabolism(14) and also promotes the production of amyloid plaques in rodents, a hallmark of Alzheimer’s disease(15, 16). Although hedonic sugar overconsumption can lead to an increased risk of metabolic and neurological diseases, including BED, a better understanding of the etiology and underlying molecular mechanisms driving binge-like eating and associated behavioral deficits such as compulsivity and anxiety is critical for the development of effective therapeutics for BED.

The prefrontal cortex (PFC) controls food-related behaviors, particularly reward processing, impulse control, and decision-making. For example, the orbitofrontal cortex (OFC) determines the hedonic (pleasure) value of food in rodents(17). The Ventromedial PFC (vmPFC) helps in decision-making based on reward and risk(18), such as choosing between a salad and a slice of cake. A recent study has shown that the PFC-lateral hypothalamus circuit controls stress-driven food intake in mice(19). Based on these studies, the PFC is a critical brain region to explore for understanding biological mechanisms of food rewards, compulsive eating, and eating disorders.

The ketogenic diet (KD) has grown in popularity as a therapeutic diet because scientific evidence supports its benefits for longevity and health span, particularly metabolic and mental health(20–22). A typical KD has a lipid content of up to 80%, adequate protein, and low carbohydrate levels, with energy produced through the catabolism of fatty acids and ketone bodies, with the brain relying heavily on ketone bodies when starved of carbohydrates(23, 24). Eating a high-energy KD is paradoxically similar to starving or restricting calorie intake, both of which rely on the mobilization of fat stores when carbohydrate availability is limited. In both cases, β-hydroxybutyrate (BHB) made from fat is a fuel molecule for the brain rather than glucose(25). In addition to being clinically beneficial in reducing epileptogenic seizures in patients with pharmacologically resistant seizures(26, 27), KD is also recommended for the treatment of glucose transporter 1 (GLUT1) deficiency syndromes(28, 29) and mood disorders(30, 31). The effects of a KD are complex, with sometimes contradictory results observed. For example, KD exacerbates neurodegeneration in mice with mitochondrial DNA toxicity in the forebrain(32) and induces p53-dependent cellular senescence in multiple organs(33).

In our previous study, we developed a clinically relevant pre-clinical model of hedonic sugar overeating and sugar dependence(34). Using the same paradigm, the major goal of this study was to investigate the underlying molecular mechanisms, specifically the synaptic, neuroimmune, and neurometabolic components of the PFC that underpin sucrose bingeing, compulsivity, and anxiety-like behavior. Furthermore, this study also investigated the therapeutic effects of nutritional ketosis on these behavioral outcomes and the molecular mechanisms underlying these effects, as KD may have therapeutic implications in cases where metabolic imbalance is associated with an increased risk and severity of a specific disease, such as BED(35, 36).

## Materials and methods

### Animals

C57BL/6J male and female mice, 6 – 8 weeks of age, were used for this study. Mice were procured from the small animal experimentation facility at the Indian Institute of Science Education and Research (IISER), Mohali, India, and housed in the Animal Experimentation facility at the BRIC-National Agri-Food and Biomanufacturing Institute (NABI), Mohali, India. Mice were maintained in a pathogen-free environment at 25 ± 2°C on a 12-h light/dark cycle with food and water *ad libitum* per the NABI Animal Care and Use Committee. All mice were housed in groups of three to five mice per cage and randomly assigned to treatment conditions after 1 week of acclimatization. Experiments were conducted according to the Committee for Control and Supervision on Experiments on Animals (CCSEA) guidelines on the use and care of experimental animals. The experimental protocol was approved by the Institutional Animal Ethics Committee (IAEC) of the BRIC-NABI (**NABI/2039/CCSEA/IAEC/2023/06**). All the experiments performed in the study were blinded at every stage of analysis, and the experimenter was unaware of the treatment each animal received.

### Diets

Top Class Pvt Ltd., India, formulated the normal chow diet (CD) and the ketogenic diet (KD) using quality-matched macro and micronutrients, as previously reported(37). The control diet had ∼10% protein, 80% carbohydrates, and 10% fat as a percentage of total calories. The KD consisted of 10% protein, 0% carbohydrates, and 90% fat. The composition of the diets is described in **Table 1**.

### Two-bottle sucrose choice paradigm

The study used a two-bottle sucrose choice paradigm to induce hedonic sucrose drinking and sugar dependence (34, 38), mimicking the human behavior of consuming high-caloric desserts after a satisfactory meal. Briefly, control mice were given two bottles (bottle 1 and bottle 2) of water and CD *ad libitum*. Mice in the sucrose bingeing group were given CD and 12 h free access to one bottle each of water (bottle 1) and 10% (w/v) sucrose (bottle 2) daily during the dark phase for eight weeks. The amount of water and 10% sucrose consumed by each animal was measured on 7, 14, 21, 28, 35, 42, 49, and 56^th^ days of the experiment. The preference given by the animals for bottle 2 containing sucrose was calculated by the formula: Bottle 2 preference (%) = Liquid consumed from bottle 2 / Total liquid consumed from bottle 1 + bottle 2 *100. To avoid place preference bias in animals, the location of bottles was switched with each other on every second day of the study.

To confirm a hedonic binge episode, animals were deprived of water/sucrose for 12 hours and reinstated to a 10% sucrose solution for 2 hours. The amount of sucrose solution consumed within the given time was measured.

After eight weeks of hedonic sucrose bingeing, animals in the dietary intervention group were given KD along with exogenous BHB (Sigma-Aldrich Co., St. Louis, Missouri, USA, Cat. No. H6501) in drinking water for the next three weeks to induce nutritional ketosis, whereas their control counterparts received CD and water. BHB was supplied in drinking water at a final concentration of 0.01875 g/ml, with a daily intake aim of roughly 3 g/kg body weight. The BHB dose was calculated using prior studies that demonstrated successful ketosis induction without negative effects(39, 40).

### Blood glucose level estimation

Blood glucose levels were estimated by using a commercially available glucometer and disposable glucose test strips (AccSure, Blood glucose monitoring system, TD-4183).

### Ketone bodies estimation

β-hydroxybutyrate (BHB) levels were estimated by using a commercially available kit according to the instructions provided by the manufacturer (Sigma-Aldrich, St. Louis, MO, USA; Cat# MAK134). In brief, BHB levels were determined using an enzymatic assay based on 3-hydroxybutyrate dehydrogenase-catalysed reactions, in which the change in nicotinamide-adenine dinucleotide (NAD^+^/NADH) absorbance measured at 340nm was directly related to BHB concentrations.

### Open field test

The open field test (OFT) was used to assess anxiety-like behavior in mice. This test is based on the natural aversion of animals to exploring new areas, where anxiety levels are inversely proportional to the preference of rodents to stay in the central arena. The experimental setup consisted of an arena (50 cm × 50 cm × 40.5 cm) placed at a sufficient height to prevent animals from escaping. The arena was divided into central and peripheral zones. The mice were placed in the central zone, and the activity of each animal was recorded for 15 minutes using ANY-maze™ tracking software (Stoelting Co., Wooddale, IL, USA). Test chambers were wiped with 70% (v/v) ethanol in between tests to remove any previous scent cues. The ethanol was allowed to dry completely before testing. Parameters such as total distance travelled, freezing time, and time spent in the central zone were measured.

### Marble-burying test

The marble-burying test (MBT) was used to assess compulsive-like behavior in mice(41). Mice were placed individually in small cages (29.0 cm x 17.5 cm), in which 20 marbles were equally distributed on top of mouse bedding (5 cm deep). A lid was placed on top of the cage to prevent the mouse from jumping out of the cage during the test. Mice were left undisturbed for 30 minutes under low light conditions, after which the number of buried marbles (those covered by bedding three-quarters or more) was counted by an observer blinded to experimental conditions.

### Enzyme-linked immunosorbent assay

Inflammatory cytokines were measured by enzyme-linked immunosorbent assay (ELISA) in the PFC according to the manufacturer’s standard protocol (Elabscience, Houston, TX, USA). Briefly, 20 mg of PFC tissue was homogenized in phosphate buffer saline (PBS) and then sonicated for 5 minutes. The homogenate was centrifuged at 5000 X g for 10 minutes at 4°C, and the supernatant was then collected. 100µl of supernatant and standards were added to a pre-coated 96-well ELISA plate and incubated at 37°C for 90 minutes. After incubation, the solution was aspirated, and each well received 100μL of biotinylated antibody detection working solution (containing biotinylated anti-mouse IL-1β, IL-10, and IL-6 antibodies) and was incubated for 1 hour. After incubation, the solution was aspirated, and wells were washed three times with wash buffer (300 μL/well) at one-minute intervals. Next, 100 μL of a secondary horseradish peroxidase (HRP) conjugated antibody was added to each well and incubated for 30 minutes at 37°C. After incubation, the solution was aspirated, and the wells were washed five times with wash buffer. After adding 90 μL substrate solution to each well, the plate was incubated for 30 minutes in the dark. After incubation, 50 μL of stop solution was added, and absorbance was measured at 450nm using a microplate spectrophotometer (SpectraMax i3x, Molecular Devices, San Jose, CA, USA).

### Isolation of cerebral microvessels

Brain microvessels were isolated from the cortical region of male and female mice as described previously(42). The animals were euthanized, and the brain tissue was carefully transferred to a petri dish with precooled MCDB 131 medium (Sigma-Aldrich Co., St. Louis, Missouri, USA). Cortical tissue was carefully transferred and homogenized in a loose-fitting, 2-ml Dounce tissue grinder in 1 ml of MCDB 131 medium at a steady pace, starting with ten strokes and not rotating the pestle. Next, the homogenized tissue was transferred to 15 ml tubes, and 7 ml of MCDB 131 medium was added to the homogenate, yielding a total volume of 8 ml. The contents were then centrifuged at 2,000 x g for 5 minutes at 4°C, the supernatant was discarded, and the tubes were inverted on a paper towel to absorb any excess medium or debris that clung to the sides. The resulting pellets were initially resuspended in 1 ml of 15% (w/v) dextran-DPBS solution. To equally dissociate the pellet, add 1 ml of 15% (w/v) dextran solution and carefully dissolve with a Pasteur pipette. Then, 7 ml of 15% (w/v) dextran solution was added to reach the final volume of 8 ml. Next, the solution was centrifuged at 10,000 x g for 15 minutes at 4°C. The pellet was dissolved in 1 ml of DPBS, filtered through a 45-μm cell strainer, and washed with <10 ml of DPBS. Vessels on the top of the filter were removed by inverting the filter and washing with 10-20 ml of MCDB 131 media containing 0.5% (w/v) BSA. Microvessels isolated using this approach ranged in diameter from 5 µm to 40 µm. They could be classified as precapillary arterioles, capillaries, or venules based on their size and marker expression. For RNA isolation, solutions were simply centrifuged at 5,000 x g for 10 minutes at 4°C to produce the final microvessel pellet, which was then processed according to the RNA isolation methodology.

### Blood-brain barrier permeability test

To evaluate blood-brain barrier (BBB) permeability in various brain regions of male mice, the animals were intraperitoneally injected with 10% (w/v) sodium fluorescein (NaF, Sigma-Aldrich, St. Louis, MO, USA) dissolved in normal saline and allowed to circulate for 45 minutes. The animals were then anesthetized with 1.5% isoflurane before blood was drawn retro-orbitally. Furthermore, the animals were transcardially perfused with ice-cold PBS. The brain tissues were extracted, weighed, and homogenized in 1 ml PBS. The homogenate was centrifuged at 1250 x g for 5 minutes, then the supernatant was collected. The supernatant was mixed with 20% (w/v) trichloroacetic acid (TCA) in a 1:10 ratio and incubated at 4°C for 24 hours to remove background fluorescence. The serum samples were diluted with 20% TCA. After incubation, the solution was centrifuged at 10,000 x g for 15 minutes to remove any precipitated protein. The supernatant was then diluted with an equal volume of borate buffer (0.05M, pH 10) to achieve a final TCA concentration of 10%. The fluorescence was measured using a 96-well plate spectrophotometer (SpectraMax i3x, Molecular Devices, San Jose, CA, USA) with an excitation wavelength of 440 nm and an emission wavelength of 520 nm. A standard curve (0-500ng) was used to determine the concentration of NaF in serum and brain tissue. The results were shown as NaF diffusion percentage of the control.

### Quantitative PCR

Animals were sacrificed by cervical dislocation. The brain was dissected, prefrontal cortex (PFC) and hippocampus regions were harvested and stored in RNA later (Sigma-Aldrich, St. Louis, MO, USA). Total RNA was isolated using a TRIzol reagent (BR Biochem Life Sciences Pvt Ltd) as per the manufacturer’s instructions. cDNA was prepared using RevertAid First Strand cDNA Synthesis Kit (Thermo Fischer Scientific, Waltham, MA, USA) by using 1 μg of total RNA. Primers were designed by using the Primer3Plus primer designing tool(43). qPCR reactions were assembled using synthesized cDNA, SYBR Green master mix (Himedia Pvt Ltd, Maharashtra, India) and 100 nM primers (Eurofins Genomics India Pvt Ltd, Bangalore, India) diluted to 5 nM final concentration. Relative mRNA expression was calculated using the 2^−ΔΔCT^ method(44). Beta-Actin *(Actb)* and Hypoxanthine-guanine phosphoribosyltransferase (*Hprt*) were used as reference genes for the expression analysis of genes of interest. The list of primers used in the study is provided in **Table 2**.

### Statistical analyses

All statistical analyses were performed in GraphPad Prism 8.1.2 (GraphPad Software, La Jolla, CA, USA). The normality assumption was verified with the Shapiro-Wilk test. Results are presented as mean ± SEM unless otherwise stated. Statistical significance was calculated by an unpaired t-test for group-wise comparisons unless otherwise stated. Statistical differences between multiple groups were determined using two-way ANOVA with sex and diet as the primary variables, followed by the Bonferroni post hoc test. A Pearson correlation analysis was performed to examine the relationship between mRNA levels and behavioral phenotype. Outliers were checked for all the treatment groups during the development of graphs.

## Results

### 1. Sucrose bingeing for eight weeks induces hyperglycemia without affecting the body weight of male and female mice

Male and female mice in the sucrose bingeing group preferred sucrose solution over water when given food *ad libitum* and a free choice between water and 10% sucrose solution for eight weeks and consumed more liquid from bottle 2 (sucrose) than their water controls throughout the experiment **(Figure 1B – 1E)**. No difference in total food intake (data not shown) and body weight was observed in male **(Figure 1F)** and female **(Figure 1G)** mice over eight weeks of sucrose bingeing. Female mice consumed more sucrose than male mice throughout the study (F (1, 80) = 67.03; p<0.0001; **Figure 1H**). Male (p=0.0381) and female (p=0.0138) mice in the sucrose bingeing group had higher blood glucose levels than their water controls (F (1, 36) = 14.16; p=0.0006; **Figure 1I**) and consumed more sucrose during 2 h of sucrose reinstatement after 12 h of sucrose deprivation (F (1, 16) = 40.63; p<0.0001; **Figure 1J**). Overall, these findings indicate that when given the choice between 10% sucrose and water, both male and female mice prefer sucrose solution over water and consume more sucrose. Furthermore, female mice drink more sucrose solution than males. Sucrose bingeing for eight weeks induces hyperglycemia and binge-like episodes in male and female mice without affecting their body weights.

**Figure 1.**
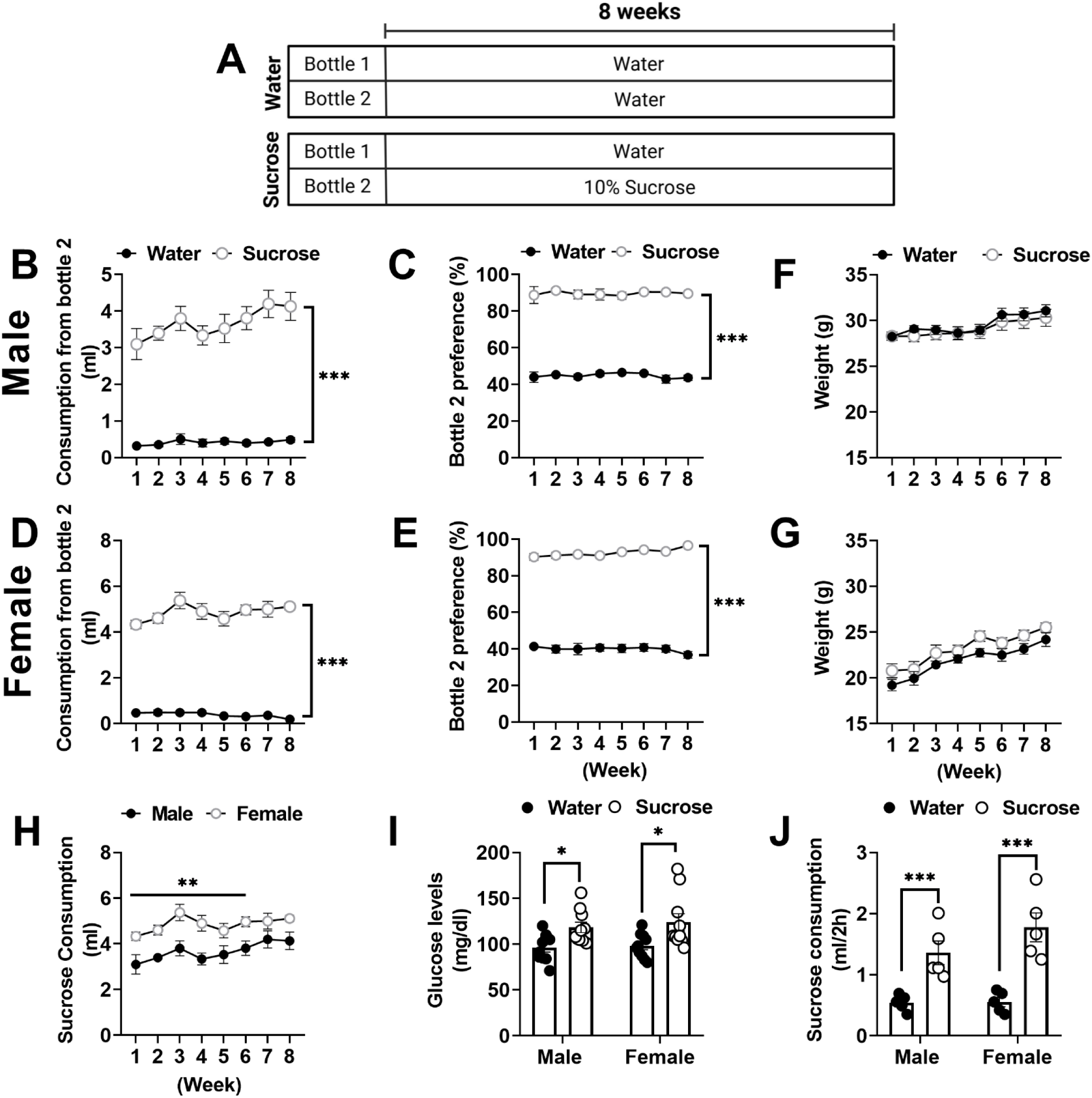
Schematic diagram of the two-bottle sucrose choice paradigm followed for the study. **(A)**. Time course analysis of 8 weeks of sucrose consumption and sucrose preference in males **(B, C)** and females **(D, E)**. Time course analysis of body weight of male **(F)** and female **(G)** mice. Time course analysis of 8 weeks of sucrose consumption and preference in male vs female **(H)**. Bar graph representing the blood glucose level in male and female mice after 8 weeks of sucrose bingeing **(I)**. Bar graph showing sucrose consumed during 2 h of sucrose reinstatement after 12 h of sucrose deprivation **(J)**. Data was analysed using two-way ANOVA with a Bonferroni post hoc test. (\**p*<0.05, \*\**p*<0.005, \*\*\**p*<0.0005; n=10-12 per treatment group).

### 2. Sucrose bingeing induces anxiety-like and compulsive-like behavior in male and female mice

OFT and MBT were used to evaluate diet effects on locomotor activity, anxiety-like and compulsive-like behavior in both male and female mice. After eight weeks of sucrose bingeing, male mice spent less time in the center zone (p=0.0145; **Figure 2A**) and had a longer freezing time (p=0.0423; **Figure 2B**) than their water counterparts. A similar trend was observed in sucrose-binged female mice, where females spent significantly less time in the center zone (p=0.0053; **Figure 2D**) and had a longer freezing time (p=0.0112; **Figure 2E**) than their water control counterparts, indicating an anxiogenic behavior. Furthermore, sucrose bingeing for eight weeks showed lower locomotor activity in both male (p=0.0002; **Figure 2C**) and female (p=0.0189; **Figure 2F)** mice as indicated by shorter distances moved in the open field arena compared to their control counterparts.

**Figure 2.**
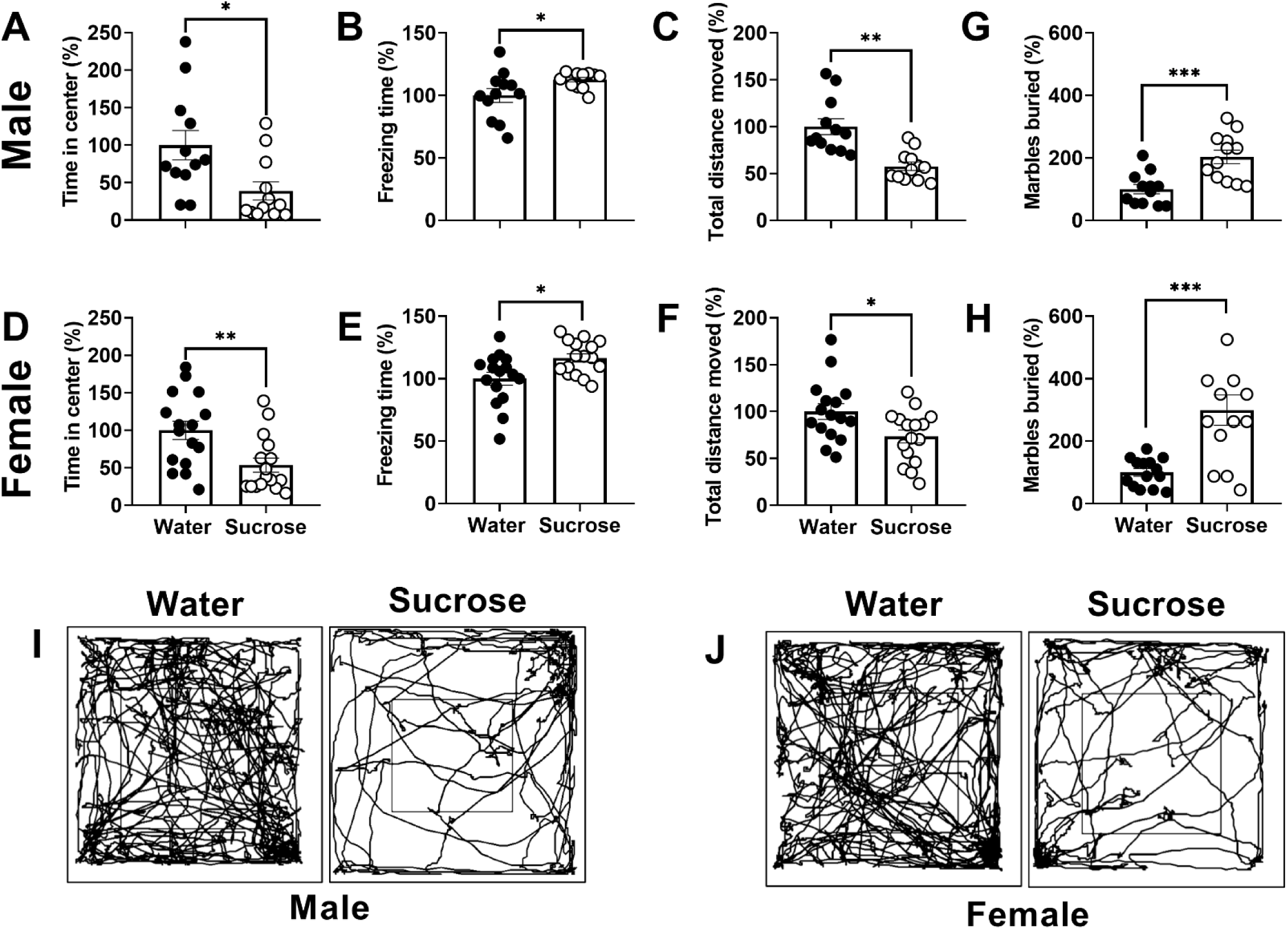
Sucrose bingeing induces compulsive-like and anxiety-like behavior in male and female mice: After eight weeks of sucrose bingeing, both male and female mice were subjected to OFT and MBT to assess compulsive-like and anxiety-like behaviors. Bar graph showing time spent in the center zone **(A, D)**, freezing time **(B, E)** and total distance moved **(C, F)** by male and female mice in the OFT. Bar graph showing the number of marbles buried in the MBT by males **(G)** and females **(H)**. Track plots of male **(I)** and female **(J)** mice in the OFT. Data was analysed using an unpaired t-test. Compare to water control *\*p*<0.05, \*\**p*< 0.005, *\*\*\*p* < 0.0005; n=10-12 per treatment group.

MBT was further used to assess the compulsive-like behavior in sucrose-binged male and female mice and to confirm that the anxiety-like symptoms observed in sucrose-binged males and females in the OFT were not due to changes in the animals’ locomotor activity. In the MBT, both male (p=0.0007; **Figure 2G**) and female (p=0.0004; **Figure 2H**) mice in the sucrose bingeing group buried significantly more marbles than their control counterparts, indicating that sucrose bingeing induces anxiety-like and compulsive-like behavior. Overall, our findings show that sucrose bingeing for eight weeks induces negative mood effects, such as anxiety and compulsive-like behavior in both males and females.

### 3. Transcription of dopaminergic receptors and synaptic plasticity marker genes in the prefrontal cortex (PFC) correlates with sucrose bingeing, anxiety-like and compulsive-like behavior in male and female mice

Prefrontal dopamine D1 (*Drd1)* and D2 (*Drd2*) receptors regulate a variety of executive functions such as assessment and decision making on risk vs reward(45). To determine how sucrose bingeing by choice affects dopaminergic signalling in the PFC, we first examined the mRNA expression of *Drd1* and *Drd2* in the PFC of male and female mice. Transcript levels of *Drd1* (p=0.0411) and *Drd2* (p=0.0411) were significantly higher in the PFC of sucrose-binged males compared to their water controls **(Figure 3A)**, whereas mRNA expression of *Drd2* (p=0.0152) but not *Drd1* was upregulated in sucrose-binged females **(Figure 3B)**.

**Figure 3.**
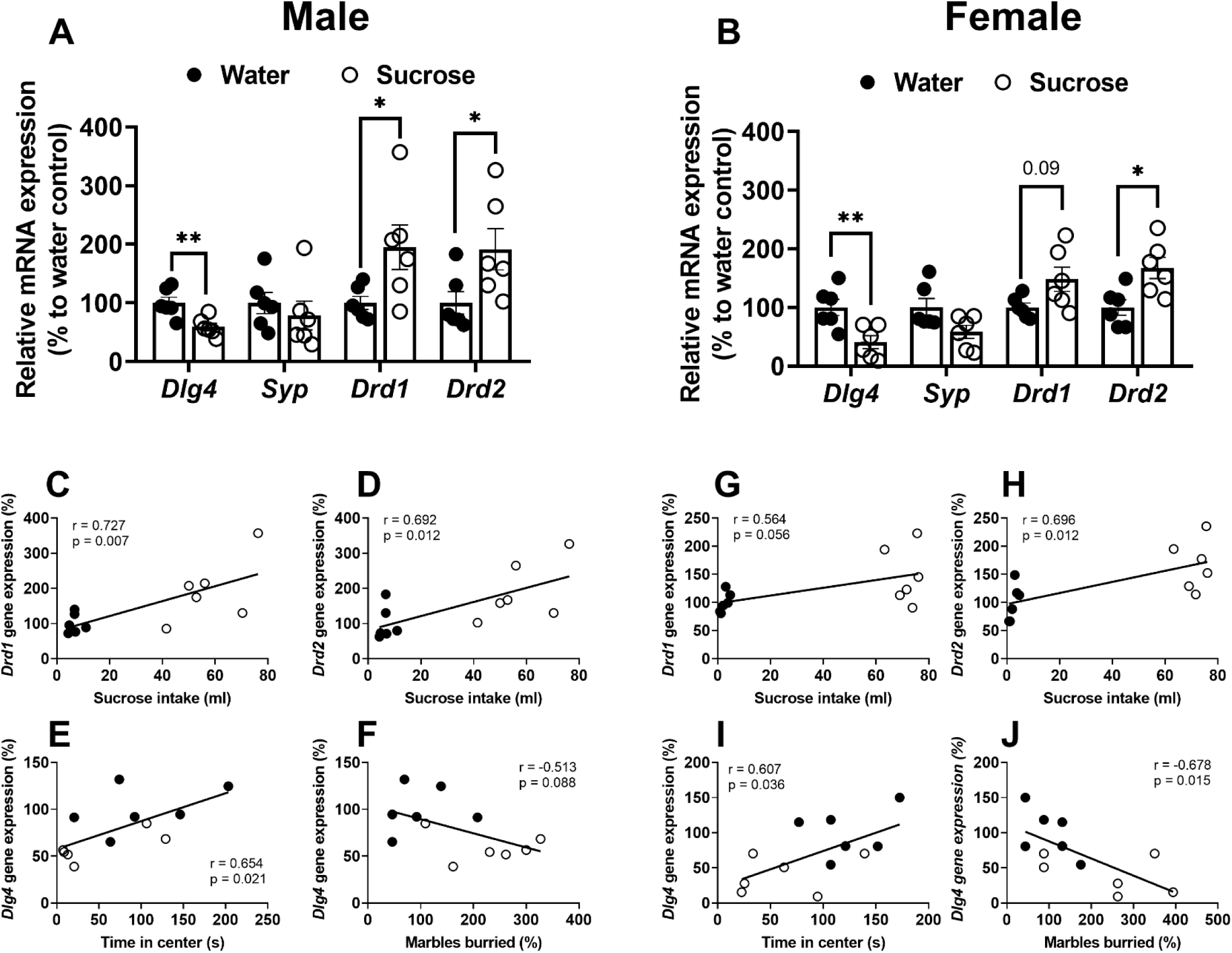
Effect of sucrose bingeing on the mRNA expression of dopaminergic receptors and synaptic plasticity markers in the PFC of male and female mice: Bar graph showing the mRNA expression of *Dlg4*, *Syp*, *Drd1* and *Drd2* in the PFC of male **(A)** and female **(B)** mice. Data was analysed using an unpaired t-test. Compared to water controls *\*p*<0.05, \*\**p*<0.005; n=6 per treatment group. Pearson’s correlation between the mRNA expression of synaptic plasticity markers and behavioral outcomes in sucrose-binged male and female mice. The correlation of *Drd1* and *Drd2* gene expression in the PFC and sucrose intake in males **(C, D)** and females **(G, H)**. The correlation of *Dlg4* mRNA levels in the PFC and anxiety parameters in males **(E, F)** and females **(G, H)**.

To further assess the impact of sucrose bingeing on synaptic plasticity, we analyzed the mRNA expression of pre-(*Syp)* and post-(*Dlg4*) synaptic markers in the PFC of male and female mice. While the mRNA expression of *Syp* in the PFC remained unchanged, the transcription of *Dlg4* was significantly downregulated in the PFC of sucrose-binged males (p=0.0087) and females (p=0.0087) compared to their water-control counterparts, indicating compromised synaptic plasticity in the PFC.

Pearson correlation analysis was performed between the transcript levels of affected neuronal markers in the PFC and their correlation with behavioral outcomes in male and female mice. The mRNA levels of *Drd1* (r=0.727; p=0.007) and *Drd2* (r=0.692; p=0.012) were positively correlated with the amount of sucrose consumed by male mice during the last day of the experiment **(Figures 3C and 3D**). However, *Dlg4* mRNA expression in the male PFC was positively correlated with the time spent in the center zone during OFT (r=0.654; p=0.021) by sucrose-binged male mice **(Figure 3E)**.

In females, there was no correlation between PFC *Drd1* mRNA expression and sucrose intake (r=0.564; p=0.056); however, *Drd2* mRNA expression was positively correlated with sucrose consumed during the last day of the study (r=0.696; p=0.012). *Dlg4* mRNA expression in the PFC of females was positively correlated with the time spent in the center zone during OFT (r=0.607; p=0.036) and negatively correlated with the total number of marbles buried during MBT (r=-0.678; p=0.015) by sucrose-binged females. Overall, these findings suggest that prolonged sucrose bingeing promotes anxiety-like and compulsive-like symptoms through modulating dopaminergic signaling and synaptic plasticity in the PFC.

### 4. Sucrose bingeing induces neuroinflammation in the PFC of male and female mice

Previously, we demonstrated that four weeks of sucrose bingeing causes a pro-inflammatory neuroimmune response in the PFC and nucleus accumbens of male and female mice, with increased IL-1β and IL-10 levels(34). To determine if neuroinflammation persists even after prolonged sucrose eating or if the previously observed elevated neuroinflammation in the brain regions was the initial neuroadaptive response to sucrose overeating, we first evaluated the mRNA expression of various pro- and anti-inflammatory marker genes (*Il1β, Il10, Ccr2, Rage*) in the PFC of sucrose binged male and female mice **(Figure 4)**.

**Figure 4.**
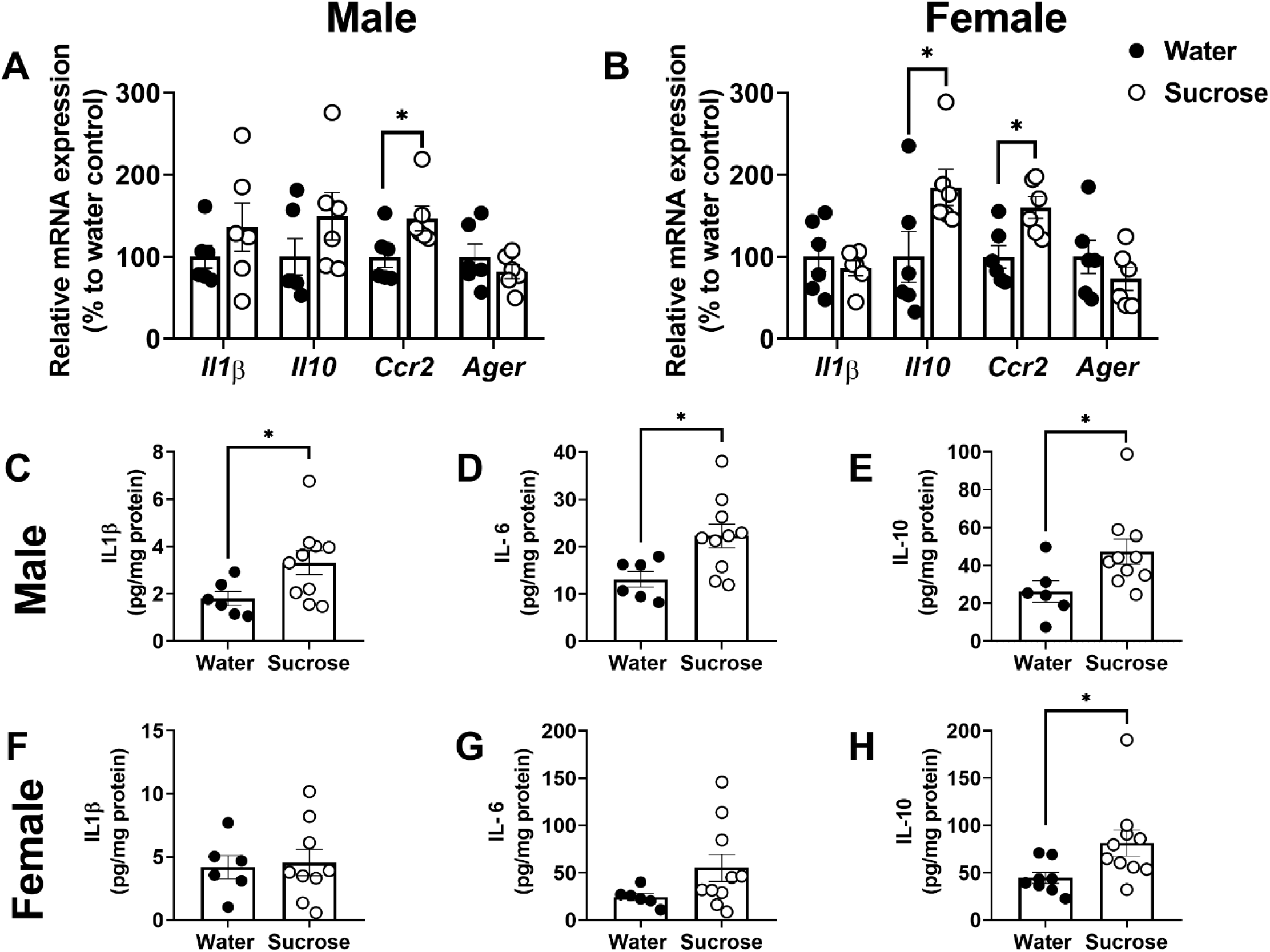
Effect of sucrose bingeing on the neuroinflammatory markers in the PFC of male and female mice: Bar graph showing the relative mRNA expression of *Il1β, Il10, Ccr2 and Ager* in the PFC of male **(A)** and female **(B)** mice. Bar graph showing the protein levels of *Il1β* **(C, F)**, *IL6* **(D, G)**, and *IL10* **(E, H)** in the PFC of male and female mice. Data was analysed using an unpaired t-test. Compared to water controls (\**p*<0.05; n=8-10 per treatment group).

Although no changes were observed in the mRNA levels of *Il1β, Il10,* and *Rage*, prolonged sucrose bingeing increased mRNA expression of *Ccr2* (p=0.0411) in the PFC of male mice compared to water controls **(Figure 4A)**. In females, sucrose bingeing increased the transcription of *Il10* (p=0.0411) and *Ccr2* (p=0.0260) genes in the PFC compared to the water control group. However, there was no change in the transcript levels of *Il1β* and *Rage* **(Figure 4B)**. The alterations in the mRNA levels of neuroinflammatory markers were region-specific, as no such changes were observed in the hippocampus of sucrose-binged male and female mice **(Figure S1)**.

We further measured the protein levels of pro- and anti-inflammatory cytokines in the PFC of male and female mice. Sucrose-binged male mice had higher levels of IL-1β (p=0.0484), IL-6 (p=0.0225) and IL-10 (p=0.0464) in the PFC compared to their water controls (**Figures 4C-4E)**. However, sucrose bingeing did not affect the levels of IL-1β and IL-6 in the PFC of the females; elevated levels of IL-10 (p=0.0389) were found in the PFC of sucrose-binged females **(Figures 4F-4H)**. These results confirm sustained neuroinflammation in the PFC of male and female mice following prolonged sucrose eating.

### 5. Sucrose bingeing compromises blood-brain barrier (BBB) integrity in male mice

We debated whether the observed neuroinflammation in this study was attributable to the compromised BBB integrity in the brain regions of male and female mice, as hyperglycemia perturbs the BBB integrity through its effects on endothelial cell features and function(46). To test this hypothesis, we first looked into how sucrose bingeing affected the mRNA expression of junctional proteins and BBB integrity markers (*Cldn5, Ocln, Tjp1, Mfsda2, Pecam*) in isolated microvessels, the smallest functional unit of the BBB from the cortex of male and female mice **(Figure 5B and 5C)**. However, sucrose bingeing did not affect the transcription of *Cldn5, Tjp1, Mfsd2a* and *Pecam*; the mRNA levels of *Ocln* was decreased (p=0.0286) in the microvessels isolated from the cortex of sucrose-binged males **(Figure 5B)**. In females, sucrose bingeing increased the transcription of *Mfsd2a* (p=0.0286) in the isolated microvessels from the cortex; however, no change was observed in the mRNA levels of *Cldn5, Ocln, Tjp1* and *Pecam* **(Figure 5C)**.

**Figure 5.**
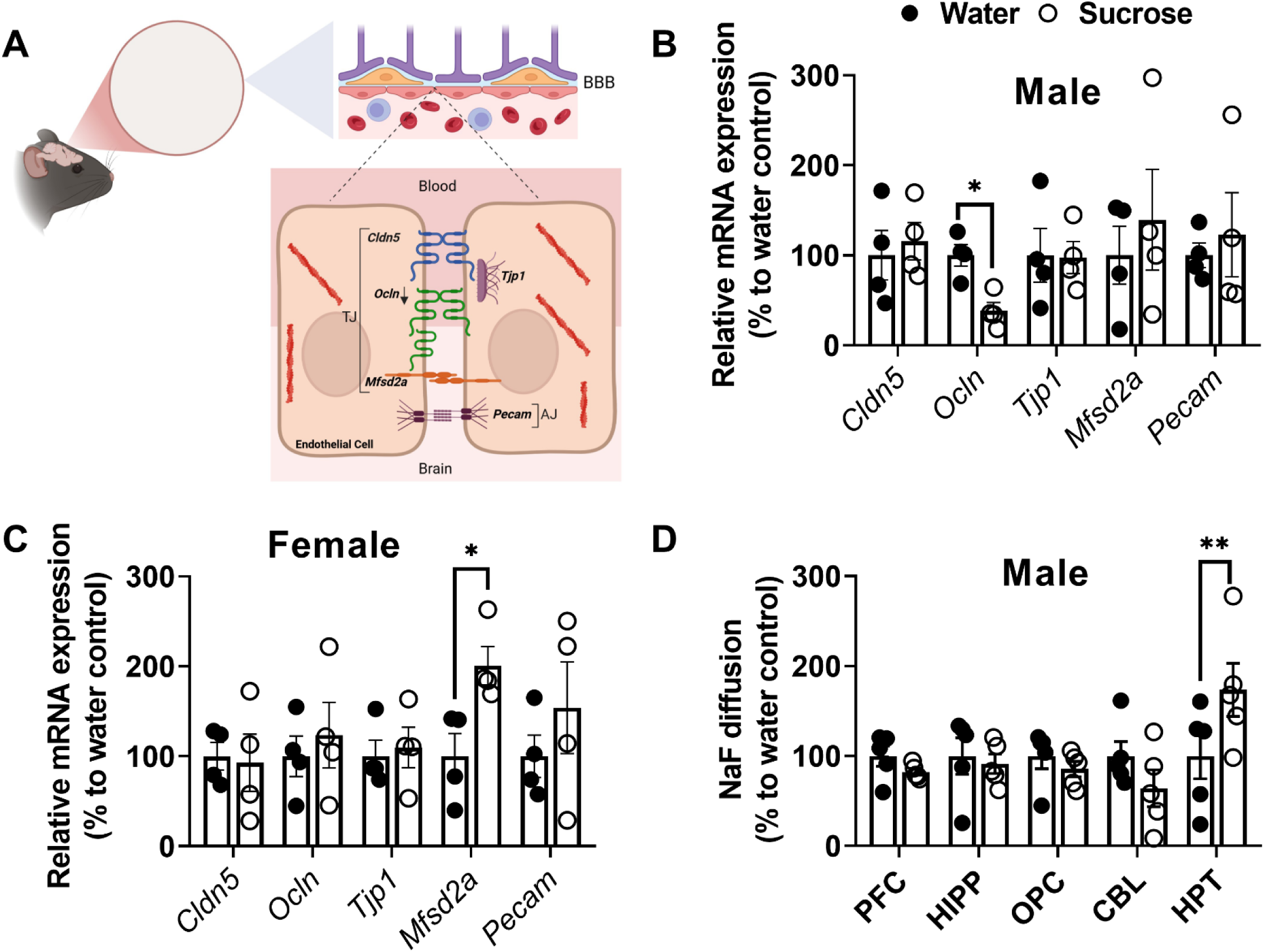
Sucrose bingeing increases region-specific blood-brain barrier (BBB) permeability in males while inducing molecular adaptation in the cortical microvessels of male and female mice: Schematic diagram showing different junctional proteins and BBB integrity markers in the microvessels, the smallest functional unit of BBB **(A)**. Bar graph showing the mRNA expression of *Cldn5, Ocln, Tjp1, Mfsda2* and *Pecam* in the isolated cortical microvessels of male **(B)** and female **(C)** mice. The BBB permeability of the prefrontal cortex, hippocampus, occipital cortex, cerebellum, and hypothalamus of male mice was assessed using the sodium fluorescein (NaF) tracer dye (molecular weight: 376 Da). Bar graph showing the percentage of NaF dye diffused in the different brain regions of male mice **(D)**. Data was analysed using a. unpaired t-test. (*p<0.05, **p<0.005; n=4-5 per treatment group).

To determine whether decreased mRNA expression of *Ocln* in males’ microvessels affects the functional properties of the BBB, we measured BBB permeability in various brain regions of males using the tracer dye sodium fluorescein (NaF; molecular weight: 376 Da; **Figure 5D**). There was no change in BBB permeability in any of the brain regions studied, including the PFC, hippocampus, occipital cortex, and cerebellum, except for the hypothalamus of sucrose-binged male mice, which showed elevated NaF levels (p=0.003) when compared to their water control counterparts **(Figure 5C)**. These findings suggest that sucrose bingeing causes molecular adaptation at the BBB site and increases BBB permeability in a sex- and brain region-specific manner, particularly in males’ hypothalamus.

### 6. Sucrose bingeing increases the mRNA expression of genes related to glucose metabolism while suppressing ketone oxidation pathway genes in the PFC of male and female mice

A high sucrose diet has been shown to induce insulin resistance with compromised energy metabolism in glial cells, impairing their phagocytic and debris removal capacity(14). Neural transmission and neuroinflammation are both energy-intensive processes, so the brain may switch its metabolism by utilizing local brain lipids and ketones as key energy sources to meet the high energy demand of neurons and neuroimmune cells for efficient functioning in sucrose-binged males and females. To confirm this hypothesis and to determine the metabolic correlates of sucrose bingeing-induced behavioral deficits, neuroinflammation and neurovascular dysfunctions, we assessed the mRNA expression of rate-limiting genes for glucose transport and metabolism (*Slc2a3, Slc2a4, Glo1*), ketogenesis and ketone oxidation (*Hmgcs2, Slc16a1, Bdh1, Oxct1, Acat1*, *CS*) in the PFC of male and female mice **(Figure 6A and 6B**). Sucrose-binged males (p=0.0411) and females (p=0.0152) exhibited a significant upregulation in the transcription of *Slc2a4*, an insulin-responsive glucose transporter in the PFC. The transcript levels of *Glo1* were higher in the PFC of sucrose-bingeing male mice (p=0.0152) compared with their control counterpart. However, no such alternation was observed in the PFC of females. While the transcription of *Hmgcs2*, a key regulatory enzyme in ketogenesis, remained unchanged in the PFC of both sexes, transcription of *Bdh1*, which catalyzes the conversion of acetoacetate to BHB and *vice versa*, was significantly upregulated in the PFC of sucrose-binged males (p=0.0104) but not in females. Conversely, the transcription of key marker genes of the ketone oxidation pathway, including *Slc16a1* (p=0.0401), *Oxct1* (p=0.0379), and *Acat1* (p=0.0401), was significantly downregulated in the PFC of sucrose-binged males. The transcription of citrate synthase (*CS*) remained unaffected in the PFC of sucrose-binged males and females compared to their control counterparts. Overall, these findings imply that sucrose bingeing induces metabolic adaptations in the PFC, such as elevated glucose metabolism and a suppressed ketone oxidation pathway, particularly in males.

**Figure 6.**
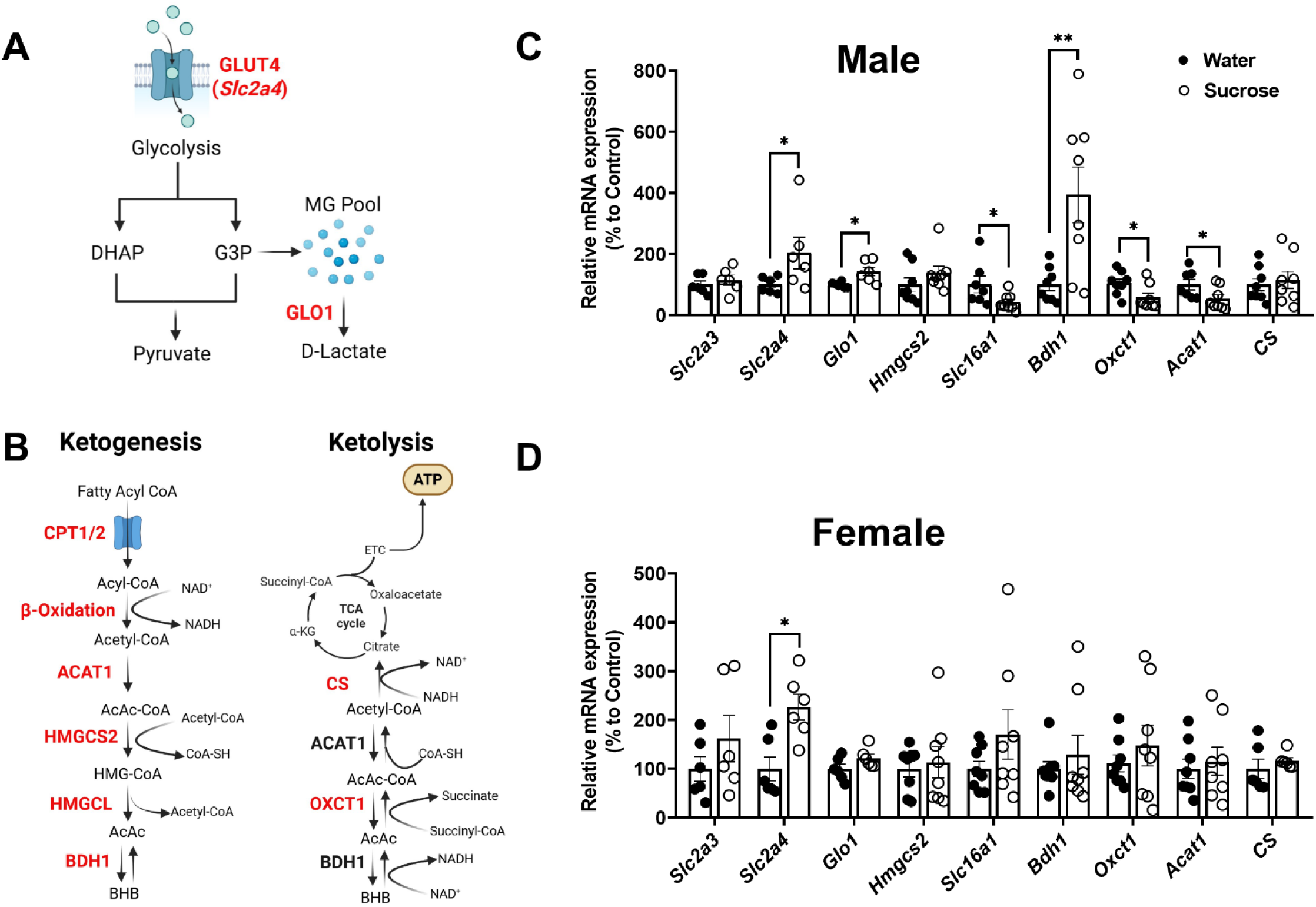
Effect of sucrose bingeing on the metabolic markers in the PFC of male and female mice: Schematics showing glucose metabolism **(A)** and mitochondrial ketone anabolic (ketogenesis) and catabolic (ketolysis) pathways **(B)** in a cell. Bar graph showing the mRNA expression of genes for glucose transport and metabolism (*Slc2a3, Slc2a4, Glo1*), ketogenesis and ketone oxidation (*Hmgcs2, Slc16a1, Bdh1, Oxct1, Acat1*, *CS*) in the PFC of male **(B)** and female **(C)** mice. Data was analysed using an unpaired t-test. (\**p*<0.05, \*\**p*<0.005; n=6-8 per treatment group).

### 7. Nutritional ketosis attenuates sucrose bingeing-induced anxiety-like and compulsive-like behavior while suppressing binge-like sucrose drinking episode in male and female mice

We further sought to determine whether the availability of alternative energy sources, such as BHB and free fatty acids, could alleviate the sucrose bingeing-induced behavioral and neurochemical deficits, as BHB bypasses glycolysis, providing a direct energy source through oxidative phosphorylation (*OXHOS*) and promoting an anti-inflammatory brain microenvironment(47). To test this hypothesis, animals in the dietary intervention group were given KD and exogenous BHB in drinking water for the next three weeks to induce nutritional ketosis following eight weeks of sucrose bingeing. In contrast, their control counterparts received CD and water. The body weight of KD-treated male (p=0.0270) **(Figures 7A)** and female (p=0.0138) **(Figures 7B)** mice significantly decreased after three weeks compared to their CD controls. Moreover, male mice showed a significant drop in blood glucose levels (p=0.0035) after three weeks of KD supplementation (F (1, 16) = 13.02; p=0.0024), while females did not show any changes in blood glucose levels **(Figure 7C)**. Male (p=0.0293) and female (p=0.0003) mice in the KD group achieved nutritional ketosis as indicated by elevated systemic BHB levels than their CD counterparts (F (1, 16) = 28.92; p<0.0001) **(Figure 7D)**.

**Figure 7.**
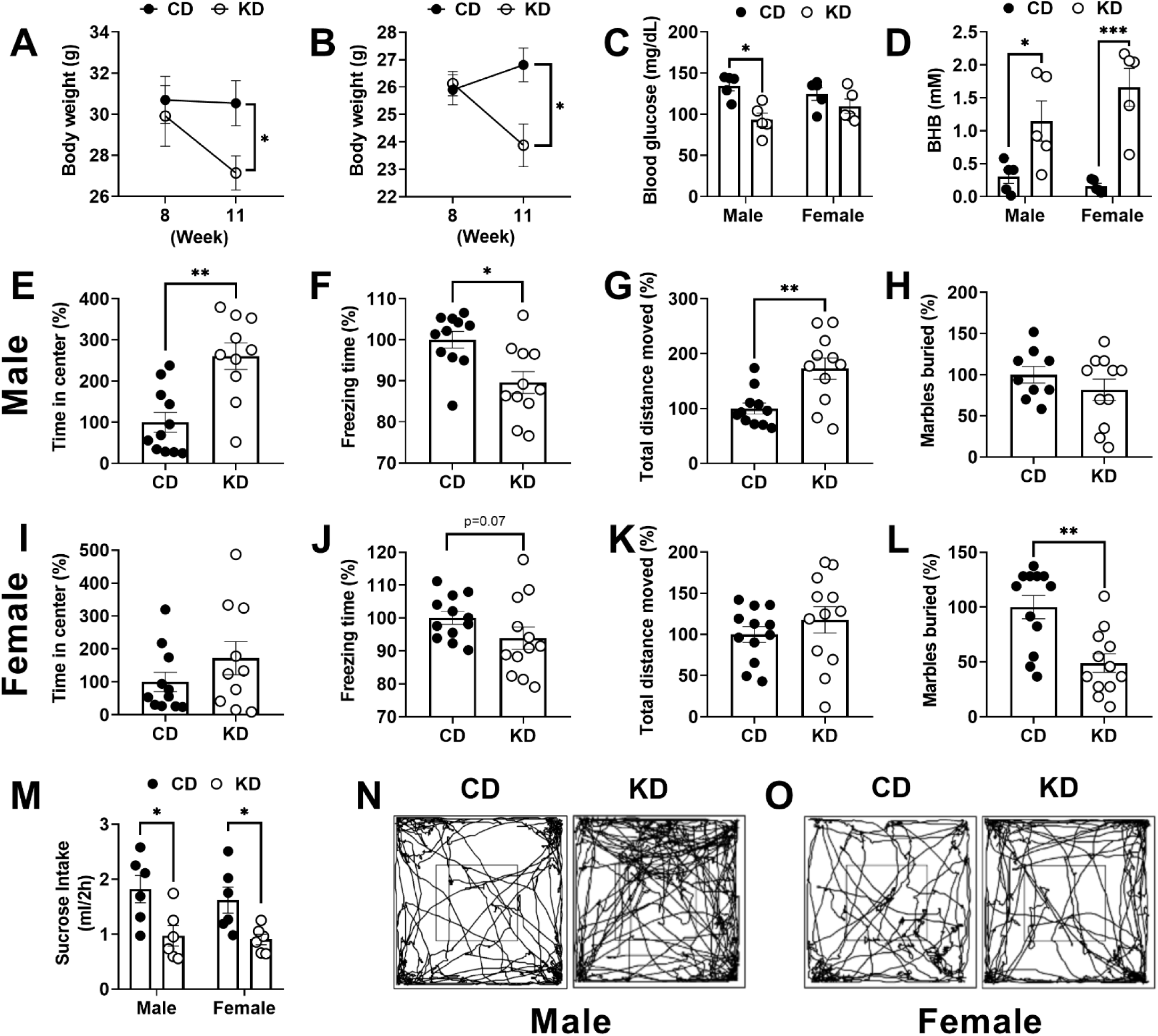
Effect of nutritional ketosis on systemic ketone levels, compulsivity, binge-like sucrose drinking and anxiety-like behavior in sucrose-binged male and female mice: After 8 weeks of sucrose bingeing, male and female mice in intervention group were supplemented with KD and BHB for 3 weeks to induce nutritional ketosis while their control counterparts received CD and water. Time course analysis of body weight in male **(A)** and female **(B)** mice. Bar graph showing the blood glucose **(C)** and BHB **(D)** levels in male and female mice. Bar graph showing time spent in the center zone **(E, I)**, freezing time **(F, J)** and total distance moved **(G, K)** by males and females in the OFT. Bar graph showing the percentage of marbles buried by males **(H)** and females **(L)** in the MBT. Sucrose intake by males and females during 2 hours of sucrose exposure **(M).** Track plots of male **(N)** and female **(O)** mice in the OFT. For body weight, blood glucose levels, BHB levels and sucrose intake, data was analysed using Two-Way ANOVA followed by post hoc Bonferroni test. For OFT and MBT, data was analysed by using an unpaired t-test. Compared to CD *\*p*<0.05, \*\*\**p* < 0.0005; n=10-12 per treatment group.

OFT and MBT were further performed to assess anxiety levels after three weeks of KD supplementation. In the OFT, time spent in the center zone was significantly (p=0.0015) higher in KD-fed males than in their CD counterparts (**Figure 7E**). Furthermore, KD supplementation reduced freezing time (p=0.0192) in males as compared to their CD controls **(Figure 7F)**. KD-supplemented male mice also travelled more distance (p=0.0083) during the OFT than their CD controls **(Figure 7G).** However, KD supplementation did not show any changes in center time, freezing time, and total distance moved during OFT in females **(Figures 7I -7K)**.

In MBT, females exposed to KD buried fewer marbles (p=0.0015) as compared to their CD control counterparts **(Figures 7L)**; however, no changes in the MBT were observed in males **(Figures 7H).** Moreover, KD supplementation significantly (F (1, 20) = 15.16; p=0.0009) reduced the sucrose intake both in males (p=0.0146) and females (p=0.0406) during sucrose reinstatement for 2 hours **(Figure 7M)**. Overall, these findings suggest that nutritional ketosis after prolonged sugar eating may promote weight loss and accelerate neurological recovery while also relieving binge-like sucrose drinking behavior in male and female mice.

### 8. Nutritional ketosis suppresses transcription of reward-related genes while increasing synaptic plasticity and anti-inflammatory markers in the PFC of sucrose-binged male and female mice

The mRNA expression of *Drd1* (p=0.0022) and *Drd2* (p=0.0152) was decreased in the PFC of KD-fed females **(Figures 8 G-H)** but not in males **(Figures 8 A-B)** as compared to their CD water controls. However, supplementation of KD significantly upregulated the transcription of *Dlg4* in the PFC of sucrose-binged male (p=0.0022) and female (p=0.0087) mice **(Figures 8C and 8I**). To further check whether KD supplementation could also attenuate the sucrose-bingeing-induced neuroinflammation, we assessed the protein levels of pro-(IL1β) and anti-inflammatory (IL-6, IL-10) cytokines in the PFC of sucrose-binged male and female mice. KD supplementation significantly increased the levels of anti-inflammatory cytokine markers such as IL-6 (p=0.0225) and IL-10 (p=0.0420) in the PFC of sucrose-binged males without affecting the IL1β levels **(Figures 8 D-F**). However, KD did not affect the cytokine levels in the PFC of sucrose-binged females **(Figures 8 K-L)**. Overall, these findings suggest that nutritional ketosis suppresses the expression of reward-related genes while promoting synaptic plasticity and an anti-inflammatory microenvironment in the PFC of males and females in a sex-dependent manner.

**Figure 8.**
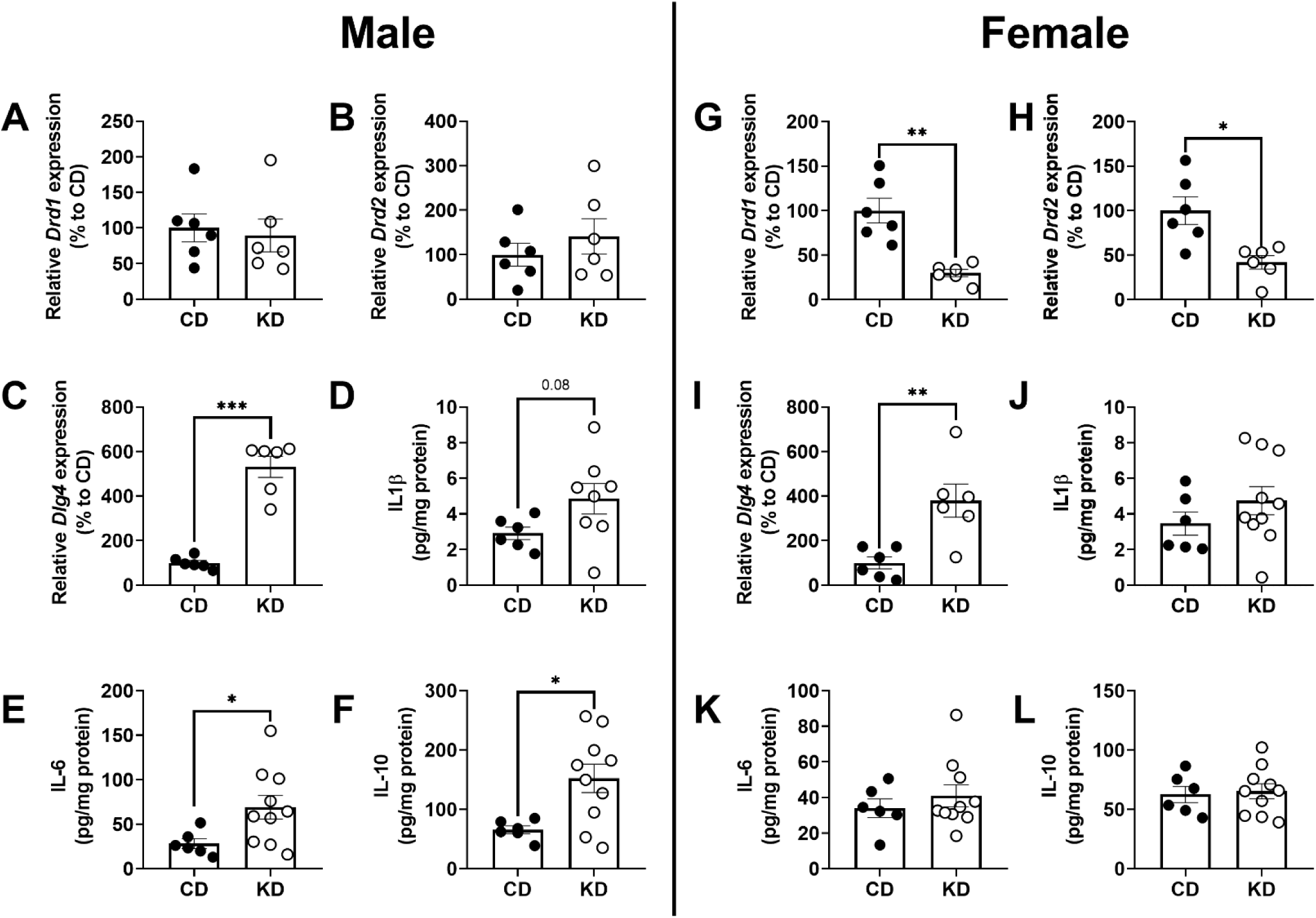
Effect of nutritional ketosis on reward, synaptic plasticity and inflammatory markers in the PFC of sucrose-binged male and female mice: Bar graph showing the relative mRNA expression of *Drd1, Drd2* and *Dlg4* in the PFC of male **(A-C)** and female **(G-I)** mice. Bar graph showing the protein levels of *Il1β* (D, J), *IL6* (E, K), and *IL10* (F, L) in the PFC of male and female mice. Data was analysed using an unpaired t-test. Compared to water controls (\**p*<0.05, \*\**p*<0.05, *\*\*\*p*< 0.0005; n=6-10 per treatment group).

**Figure 9.**
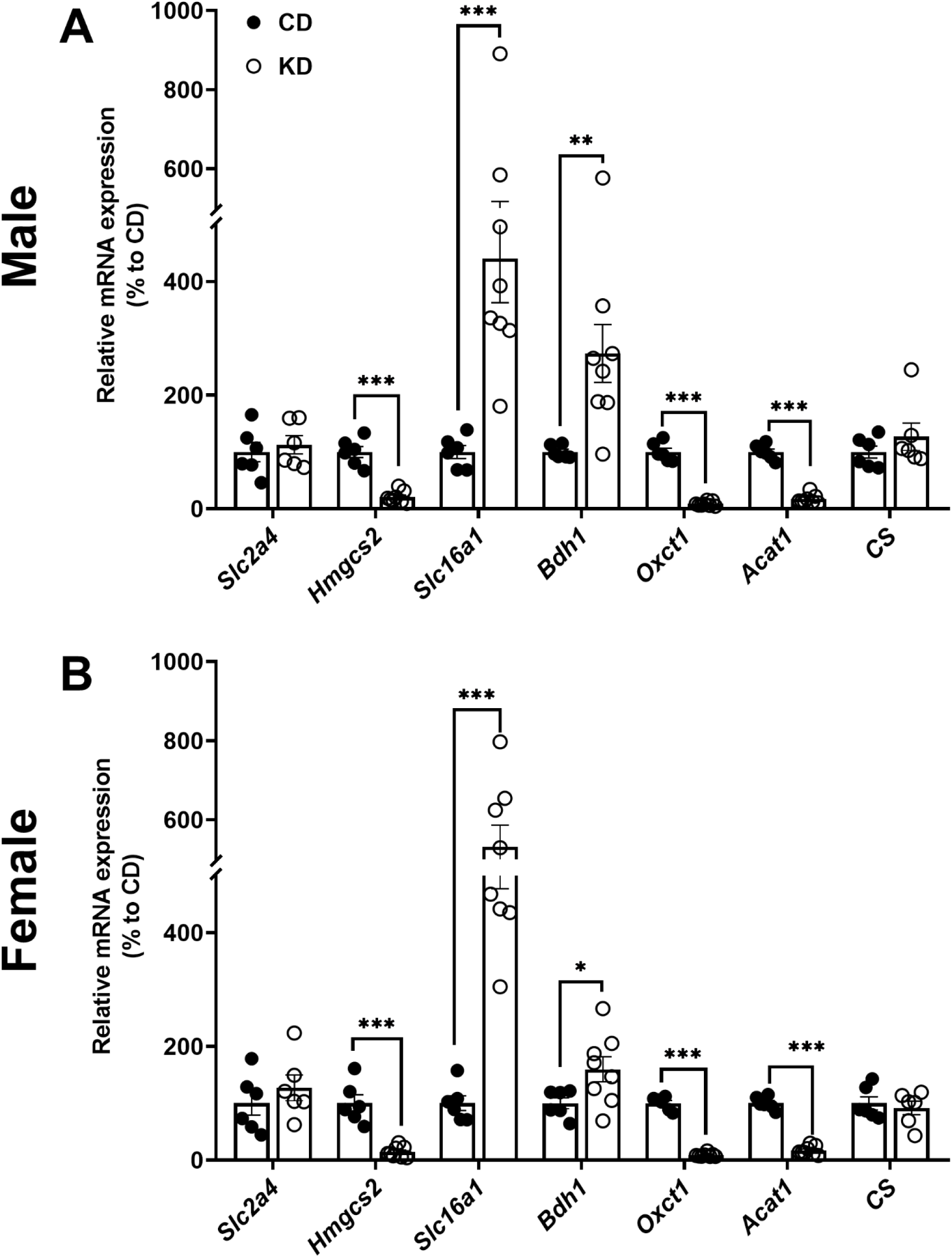
Effect of nutritional ketosis on the metabolic markers in the PFC of sucrose-binged male and female mice: Bar graph showing the mRNA expression of genes for glucose transporter (*Slc2a4*), ketogenesis and ketone oxidation (*Hmgcs2, Slc16a1, Bdh1, Oxct1, Acat1*, *CS*) in the PFC of male **(B)** and female **(C)** mice. Data was analysed using an unpaired t-test. (\**p*<0.05, \*\**p*<0.005; n=6-8 per treatment group).

### 9. Nutritional ketosis further suppresses the transcription of local ketogenesis and ketone oxidation marker genes in the PFC of sucrose-binged male and female mice

We further argued if the favorable effects of nutritional ketosis on behavioral deficits, neuroinflammation and synaptic plasticity in sucrose-binged male and female mice were attributable to its capacity to increase brain ketone oxidation in the PFC. The mRNA expression of *Slc2a4*, *Hmgcs2, Slc16a1, Bdh1, Oxct1, Acat1*, and *CS* was further analysed in the PFC of male and female mice. KD did not change the mRNA expression of *Slc2a4* in the PFC of sucrose-binged males and females compared to their control counterparts. KD decreased the transcription of *Hmgcs2,* the rate-limiting gene for ketogenesis in the PFC of sucrose-binged males (p=0.0007) and females (p=0.0007). However, KD increased the transcription of *Slc16a1 and Bdh1* but suppressed the transcription of *Oxct1 and Acat1* in the PFC of sucrose-binged males and females. However, no significant changes were observed in the mRNA expression of *CS*. These data imply that nutritional ketosis did not promote the transcription of ketone oxidation marker genes in the PFC of sucrose-binged male and female mice.

## Discussion

Although the binge-like eating of refined sugars and sugar-sweetened beverages (SSBs) is detrimental to overall health, their consumption is rapidly increasing worldwide due to their rewarding effects and may lead to the development of BED. In this study we have discovered that: 1) The preclinical model used here exhibites many of the hallmark characteristics of BED, such as sucrose bingeing in short period, compulsivity and anxiety; 2) Female mice binge more on sucrose than male subjects; 3) Sucrose bingeing for eight weeks induces anxiety and compulsive-like behavior with a concurrent increase in low grade chronic neuroinflammation, compromised region- and sex-specific BBB integrity, and altered transcription of reward, synaptic plasticity, and metabolic genes in the PFC of male and female mice; 5) Nutritional ketosis attenuates sucrose bingeing-induced behavioral deficits by improving synaptic plasticity and anti-inflammatory signalling in the PFC in a sex-dependent manner.

When mice were given chow *ad libitum* and free choice between water and 10% sucrose, both male and female mice preferred a sweet solution over water and consistently consumed more sucrose without affecting their total food intake; however, female mice consumed more sucrose than male mice, which is consistent with our previous study(48). The distinctive preference for sucrose observed in males and females may be due to biological differences between the two sexes, particularly hormonal differences. For example, estrogen has been shown to increase the detection threshold for sucrose taste in female rodents(49). Changes in females’ monthly patterns of sweet taste sensitivity and food intake caused by changes in sex hormones during the menstrual cycle have been well documented(50). These differences may influence a female’s subjective experience of sweet taste and motivation to consume sweet food and beverages. Male and female mice developed hyperglycemia following eight weeks of sucrose bingeing; however, no significant change in body weight was found in males and females, possibly because palatable food develops obesity and BED through different pathophysiological mechanisms(35). Males and females also exhibited higher sucrose consumption during two hours of sucrose reinstatement following 12 hours of sucrose deprivation. A preclinical study in rats found that prolonged sucrose bingeing decreases reward system activity by lowering neuronal firing rates in reward neurocircuitry components such as the medial prefrontal cortex (mPFC) and ventral tegmental area (VTA). The neuronal firing rates increased during active bingeing episodes of palatable meals such as sucrose, resulting in a sense of satisfaction(51), which could explain binge-like sucrose drinking during 2-hour sucrose reinstatement observed in this study. These behavioral outcomes are supported by increased mRNA expression of *Drd1* and *Drd2* receptors in the PFC of sucrose-binged males and females, as well as their correlation with sucrose drinking, because expression levels of these receptors in the PFC are associated with reward-associated responses and motivation to seek and consume hedonic substances such as sucrose(52, 53).

Prolonged sucrose bingeing significantly enhances anxiety-like and compulsive-like behaviors in both sexes. In the OFT, sucrose-binged mice spent less time in the central zone and had longer freezing periods than control animals, indicating increased anxiety-like behavior. Both male and female mice displayed this behavior, implying that sucrose bingeing may sensitize stress pathways, either through dysregulation of the hypothalamic-pituitary-adrenal (HPA) axis(54, 55) or changes in mesolimbic dopamine signaling(12), as demonstrated in our study. The MBT gave more insight into the compulsive components of behavior. Sucrose-binged mice buried substantially more marbles than controls, indicating increased compulsivity or repetitive behavior. This behavioral pattern is consistent with prior observations of distinct neurochemical responses to palatable food intake, notably in dopaminergic and cholinergic systems involved in compulsive behavior(56, 57). Reduced *Dlg4* gene expression in the PFC of sucrose-binged male and female mice and its correlation with anxiety and compulsive-like phenotype further supports the observation that high caloric diets increase the onset and progression of psychiatric disorders(7) that could be attributed to the loss of *Dlg4* by these diets, as the *Dlg4* gene is directly associated with neurological disorders(58, 59). Taken together, these findings indicate that persistent sucrose bingeing can produce a behavioral phenotype marked by increased anxiety and compulsivity. The involvement of both behavioral domains lends credence to the idea that excessive ingestion of palatable substances, including non-drug rewards such as sucrose, can activate overlapping brain processes involved in affective and compulsive disorders such as BED.

The neurobiological consequences of sucrose bingeing were explored in male and female mice, concentrating on markers of neuroinflammation and BBB integrity. Our findings show that sucrose bingeing increased neuroinflammatory responses and impaired BBB function, with unique sex-related patterns. Sucrose-binged mice had higher levels of pro- and anti-inflammatory cytokines (IL-1β, IL-6, IL-10) in the PFC compared to controls. This implies that sugar bingeing elicits a central neuroimmune response, which may further trigger metabolic abnormalities and peripheral inflammation. Previous research suggests that excessive refined sugar intake activates Toll-like receptor signaling and NF-κB pathways(60), leading to neuroinflammation. Importantly, male mice displayed higher levels of IL-1β and IL-6 in the PFC than their female counterparts may be because male mice have more cortical microglia, the primary innate immune cells of the CNS, than female mice(61), which are more sensitive to activation by any external stimuli(62).

BBB integrity was significantly impaired in sucrose-binged mice in a sex-dependent manner, as measured by mRNA expression of tight junction proteins such as claudin-5 (*Cldn5*), occludin (*Ocln*), and ZO-1 (*Tjp1*) in the isolated brain vasculature, as well as the sodium fluorescein (NaF) extravasation assay. Reduced *Ocln* mRNA expression in isolated cerebral vasculature and higher dye permeability indicated impaired BBB integrity in sucrose-binged male mice. The enhanced neuroinflammation found in this study is most likely owing to high-sucrose diet-induced BBB damage, which supports earlier findings that a high-sucrose diet causes brain angiopathy(7) and that an acute high-fat, high-sugar diet increases BBB permeability(63). Diet-induced BBB degradation may increase paracellular permeability and peripheral immune cell infiltration, aggravating neuroinflammation, as indicated by increased *Ccr2* mRNA expression in the PFC of sucrose-binged male and female mice as a proxy for invading peripheral immune cells(64). Interestingly, only males showed lower vascular *Ocln* mRNA expression and higher hypothalamic NaF levels. These regional and sex-specific effects of a high sucrose diet on BBB physiology could be attributed to differences in metabolic stress responses, vascular control, or hormonal influences on male and female endothelial cells.

Sucrose bingeing changes the metabolic profile of the PFC of male and female mice, as evidenced by increased expression of glucose metabolic genes while suppressing mitochondrial ketone oxidation marker genes in a sex-specific manner, indicating that male and female brains have distinct metabolic vulnerabilities. In contrast to previous research that a high sucrose diet causes insulin resistance in brain cells of *Drosophila melanogaster* (14), this study found that both male and female mice had increased transcription of the insulin-sensitive glucose transporter (GLUT4) encoding gene *Slc2a4* in their PFC. Furthermore, the increased mRNA expression of the *Glo1* gene in the PFC of males may be a compensatory mechanism of the brain to detoxify methylglyoxal produced by the higher glycolysis rate in the PFC of sucrose-binged males. Furthermore, sex-specific analysis revealed a decrease in the transcription of ketone oxidation pathway genes in the PFC of male mice but not in females. This could be due to differences in the regulation of important ketolysis enzymes such as succinyl-CoA:3-oxo-acid CoA-transferase 1 (SCOT; *Oxct1*) and 3-Hydroxybutyrate dehydrogenase 1 (BDH1; *Bdh1*) by the age and sex of the rodents(65), as well as sex differences in brain mitochondrial efficiency and substrate utilization, with female’s brain mitochondria having higher respiration capacity than their male counterparts(66). This impairment suggests a metabolic shift away from efficient ketone utilization by the brain exposed to a high-sucrose diet, especially in males, which may compromise energy homeostasis of brain cells during transient glucose scarcity during the sugar withdrawal phase and may lead to sugar cravings and withdrawal symptoms as observed in our previous study(34).

The current study provides compelling evidence that nutritional ketosis significantly reduces binge-like phenotype, compulsivity and anxiety-like behavior in sucrose-dependent male and female mice. The protective effects appear to result from suppressed dopamine signalling, increased anti-inflammatory microenvironment and improved synaptic plasticity in the PFC in a sex-dependent manner. Behaviorally, KD supplementation significantly reduced binge-like sucrose drinking, the number of marbles buried, and center-avoidant behavior during the OFT in sucrose-dependent male and female mice. Mechanistically, KD suppressed the transcription of *Drd1* and *Drd2* genes in the PFC of sucrose-binged female mice, indicating a blunted reward-driven neuroplasticity in females. In contrast, KD increased the levels of anti-inflammatory cytokines such as IL-6 and IL-10 in the PFC of males, not in females. Increased mRNA expression of *Dlg4* in the PFC of KD-fed males and females indicates that nutritional ketosis promotes synaptic resilience. These adjustments are most likely responsible for the observed reductions in compulsive and anxiety-like behaviors in sucrose-binged males and females. Our findings are consistent with prior research in which KD was proven to reduce drug reward, such as alcohol intake in mice as well as alcohol withdrawal symptoms in both rodents and humans(67, 68). Immunomodulation by the KD has been proposed as a therapeutic approach for treating drug addiction(69). Importantly, our findings show sex-specific effects of nutritional ketosis in neuronal, neuroimmune, and behavioral responses in sucrose-binged male and female mice, with males potentially benefiting more from KD’s anxiolytic and anti-inflammatory effects and females demonstrating more reward-circuit stabilization, emphasizing the importance of gender-specific approaches in dietary therapies for the treatment of BED.

## Conclusions

This study shows that prolonged dietary refined sugar has far-reaching consequences on the metabolism, synaptic plasticity, neuroimmune and neurovascular functions of the brain components of reward neurocircuitry, such as the PFC. These neurochemical changes may underlie the refined sugars and sugar-enriched palatable food-driven bingeing, anxiety-like and compulsive-like behaviors. The sex-specific neuroimmune, neurovascular and metabolic patterns observed in the PFC of sucrose-binged male and female mice may contribute to developing brain-targeted precision therapeutics for binge-like eating disorders. Furthermore, this study supports nutritional ketosis as a precision metabolic therapy for binge-like eating and associated affective disorders by suppressing reward-related gene expression and inducing an anti-inflammatory microenvironment in the PFC. However, additional research is needed to confirm the therapeutic efficacy of nutritional ketosis, including ketone metabolism-targeted medicines for the treatment of binge-like eating disorder in clinics.

## Compliance with Ethical Standards

### Statements and Declarations

### Funding

This work was supported by the Department of Biotechnology, Govt. of India, in the form of MK Bhan Young Researcher Fellowship and a research CORE grant from the National Agri-Food and Biomanufacturing Institute (BRIC-NABI) to MK.

### Competing Interests

Authors have no competing financial interests.

### Data availability statement

The data supporting this article have been included as part of the Supplementary Information.

### Authors’ contributions

MK obtained funding for the study. MK and CG designed all the experiments. CG performed experiments. CG and MK wrote the first draft of the manuscript. All authors edited subsequent versions and approved the final version of the manuscript.

### Ethics approval

Institutional Animal Ethics Committee (IAEC) of BRIC-NABI approved experimental protocol (NABI/2039/CPCSEA/IAEC/2023/06).

### Consent to participate

Not applicable

### Consent for publication

Not applicable

## Acknowledgements

The author thanks the Executive Director (BRIC-NABI) for the facilities. Illustrations were created by using BioRender, an online scientific image illustration tool (BioRender.com).

## Figures

**Figure S1.**
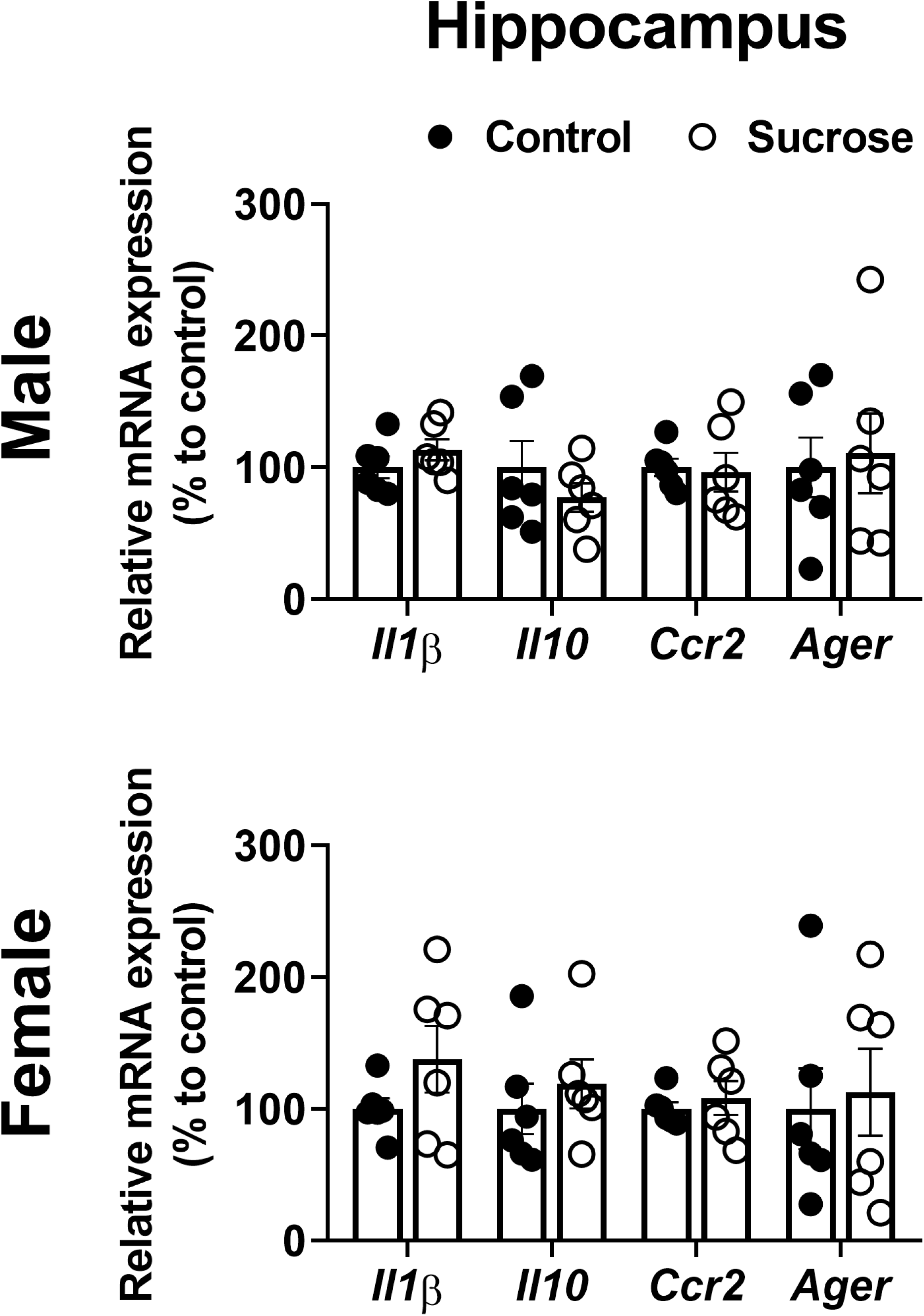
Effect of sucrose bingeing on the neuroinflammatory markers in the hippocampus of male and female mice: Bar graph showing the relative mRNA expression of *Il1β, Il10, Ccr2 and Ager* in the hippocampus of male **(A)** and female **(B)** mice.

## Notes

### Competing Interest Statement

The authors have declared no competing interest.

